# A wavelet-based approach generates quantitative, scale-free and hierarchical descriptions of 3D genome structures and new biological insights

**DOI:** 10.1101/2024.07.12.603291

**Authors:** Ryan Pellow, Josep M Comeron

## Abstract

Eukaryotes fold their genomes within nuclei in three-dimensional space, with coordinated multiscale structures including loops, topologically associating domains (TADs), and higher-order chromosome territories. This 3D organization plays essential roles in gene regulation and development, responses to physiological stress, and disease. However, current methodologies to infer these 3D structures from genomic data have limitations. These include varying outcomes depending on the resolution of the analysis and sequencing depth, qualitative results that hinder statistical comparisons, lack of insight into the frequency of the structures in samples with many genomes, and no direct inference of hierarchical structures. These shortcomings can make it difficult for the rigorous comparison of 3D properties across genomes, between experimental conditions, or species. To address these challenges, we developed a wavelet transform-based method (WaveTAD) that describes the 3D nuclear organization in a resolution-free, probabilistic, and hierarchical manner. WaveTAD generates probabilities that capture the variable frequency within samples and shows increased accuracy and sensitivity compared to current approaches. We applied WaveTAD to multiple datasets from *Drosophila*, mouse, and humans to illustrate new biological insights that our more sensitive and quantitative approach provides, such as the widespread presence of embryonic 3D organization before zygotic genome activation, the effect of multiple CTCF units on the stability of loops and TADs, and the association between gene expression and TAD structures in COVID-19 patients or sex-specific transcription in *Drosophila*.

## INTRODUCTION

The primary linear structure of genetic information within nuclei undergoes a highly specific and programmed multi-level hierarchical organization in three-dimensional (3D) space. This organization. These 3D genomic structures, including loops, topologically associated domains (TADs) and territories, facilitate interactions between distant sequences, such as enhancers and promoters, which are vital for orchestrating transcriptional programs in a developmental and tissue-specific manner (Bonev and Cavalli 2016; Szabo et al. 2019). They also position specific genomic sections in differentially accessible nuclei compartments. Alterations in these tissue-specific 3D structures have been linked to developmental abnormalities, diseases, and cancer (Ibn-Salem et al. 2014; Katainen et al. 2015; Oldridge et al. 2015; Flavahan et al. 2016; Ji et al. 2016; Valton and Dekker 2016). However, not all gene expression control is dependent on TAD and loop formation, as some genes show robust expression independent of TAD perturbations (de Laat and Duboule 2013; Williamson et al. 2019), suggesting structural as well as functional properties.

Developments in genomic methods, including several forms of high-throughput chromatin conformation capture (Hi-C) (Jerkovic and Cavalli 2021) and more recently single-cell Hi-C methods (Galitsyna and Gelfand 2021), allow generating high-resolution DNA sequence data that identify contacts between distant genomic regions. These contacts form matrices (contact matrices), which can be used to infer proximity in 3D space and predict structures such as DNA loops and TADs (Belaghzal et al. 2017), with TADs showing frequent internal interacting areas relative to loops (Schwartz and Cavalli 2017). However, the raw information used for contact maps is considerably sparse and noisy due to the limited number of informative reads per kilobase (kb) and the presence of random contacts. In the last decade, many computational and mathematical tools have been developed to infer loops and TADs based on contact matrices (Dali and Blanchette 2017; Forcato et al. 2017; Zufferey et al. 2018). All these tools integrate contact data at a given genomic scale or resolution that is decided *a priori*, often determined by the size of the genome and number of informative reads. Most studies based on human or mouse bulk Hi-C use a 50kb resolution, analyzing all the reads that connect sites within a specific 50kb genomic region with any other 50kb region across the genome. Studies in species with smaller genomes, such as *Drosophila*, tend to use finer resolutions (e.g., 10-25kb) due to the ease to obtain higher numbers of informative reads per genomic unit. The collapse of data when using broad resolutions limits the precise identification of contact locations and overlooks weaker secondary contacts within a region, while studies at finer scales generate noisier signal. These limitations are drastically intensified in single-cell studies, often requiring resolutions at or above 500kb (Ramani et al. 2017; Kim et al. 2020). Notably, all current methods to infer 3D structures based on contact matrices predict different sets of loops and TADs (number, size and locations) depending on the resolution used in the analyses, restricting biological insights from comparisons between studies and/or species. Another limitation of 3D algorithms is the difficulty in providing quantitative descriptions, further limiting accurate comparisons between structures across genomes or samples.

To address the limitations of current methodological approaches to study 3D genome organization, we developed a method and a corresponding bioinformatics pipeline to analyze contact frequencies using wavelet-transforms (WT). Proposed in the early 1900s, the popularity of WT began to take hold in the 1980s in physics and geology as a means of signal and, later, image processing (Percival and Walden 2006). When applied to genomic studies, WT can be used to study how any type of signal changes along chromosomes by decomposing it into a set of scaling and wavelet functions. These functions provide local information in both location and scale (size of the genomic region) without the need to specify the scale of interest *a priori*, making it resolution-free. Specifically, WT produces detail coefficients that represent the change in signal intensity between neighboring locations at a given scale while removing the noise and signal identified at smaller scales without loss of information. This makes it useful for extracting multi-scale signals from noisy data (Mulcahy 1996; Addison 2005; Percival and Walden 2006; Abuhamdia and Taheri 2015) and, for most genomic studies, WT would identify signals starting from the smallest possible scale—a single nucleotide. Importantly, detail coefficients are normally distributed, allowing for the assignment of independent probabilities for each denoised scale-by-location datapoint (Percival and Walden 2006) and the quantitative comparison between signals regardless of scale.

While WT has been used in genomic applications in the past, mostly with the objective of denoising data before downstream analysis (Lio 2003; Meng et al. 2013), we propose that WT is an ideal and statistically rigorous framework to analyze contact frequencies across genomes and infer loops and TADs. Moreover, we take full advantage of the resolution-free properties of WT to directly identify potential multilevel structures (e.g. loops within loops), removing the limitation of most algorithms that only identify structures adjacent to each other. Here, we describe the new method and the accompanying software package (WaveTAD) to analyze Hi-C contact data at the nucleotide level in a probabilistic, resolution-free, and hierarchical manner. We also apply WaveTAD to multiple datasets, both bulk and single-cell Hi-C, to demonstrate its benefits providing new biological insights.

## RESULTS

### Identification of loops and TADs using wavelet-transforms with WaveTAD

The use of WT for genomic analysis requires data in the form of a signal across genomes, in this case contact reads associated with a given genomic location. The WaveTAD pipeline begins by mapping paired-end Hi-C sequencing reads using BWA-MEM (Li and Durbin 2009) (**Figure 1**; see **Methods** for details). Pair reads showing intrachromosomal interactions with a minimum insert length of 500bp are then split into 5’ and 3’ groups (defining the direction of the interaction in linear space; e.g., 5’ reads interact with their paired read located 3’ along the chromosome). The coverage of each group is calculated using *genomecov* (Quinlan and Hall 2010), which transforms the data into a signal at 1-bp resolution. This signal is then subjected to a WT algorithm to identify TAD boundaries. High-value detail coefficients and corresponding probabilities generated from the wavelet-transformation allow identifying locations with a significant increase in contact frequency along a chromosome, defining TAD and loop boundaries. These boundary locations are paired with 3’ or 5’ counterparts to produce TADs that are then filtered based on the contact density at the loop anchor relative to the surrounding area (Wolff et al. 2020).

**Figure 1.**
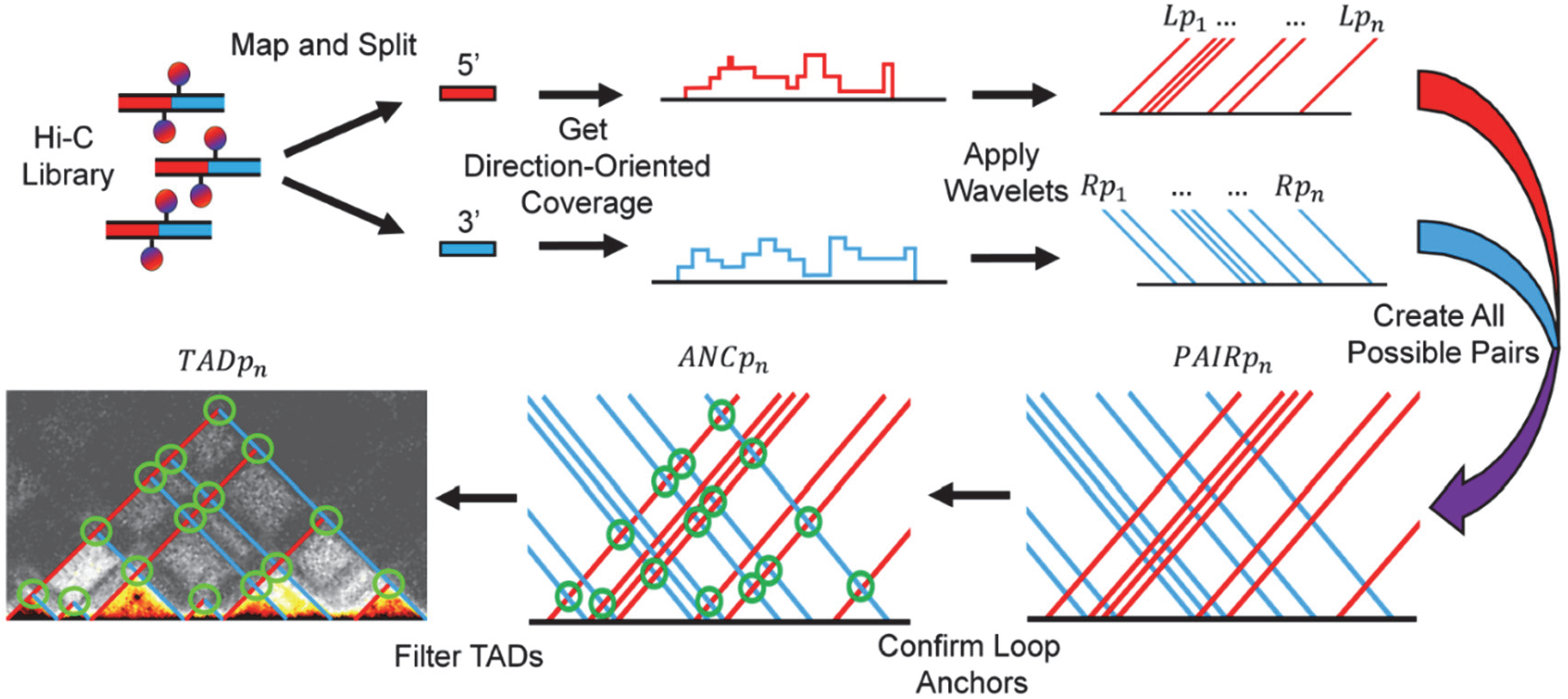
Overview of WaveTAD method. Sequencing pair-reads are first mapped and filtered for reads uniquely mapping to the reference genome, which represent valid Hi-C contacts. Based on the orientation of the mapped reads, they are split into 5’ (red) and 3’ (blue) counterparts in relation to the reference coordinates, indicating contacts downstream and upstream, respectively. Genome-wide coverage is then calculated for both the 5’ and 3’ segments and each coverage signal undergoes WT, which identifies locations in the genome displaying sharp increases in coverage and represent the left and right TAD boundaries. Importantly, this process assigns probabilities to each TAD boundary. All left-side and right-side TAD boundaries are then paired to create all possible TAD boundary pairs, which are then filtered based on the presence of a loop anchor (depicted by the green circles). Each loop anchor is assigned a probability based on the number of Hi-C contacts between the boundary pairs relative to the surrounding genomic locations. Redundant TAD calls are filtered out and the product of the probabilities of the left TAD boundary, right TAD boundary, and loop anchor are calculated and denote the probability, as a measure of stability, of the called TADs.

Considering that TAD boundaries may contain multiple architectural protein binding sites, leading to multiple location hits for the same loop anchor site, a consensus TAD boundary location is generated by applying a diamond area algorithm (Shin et al. 2016). Finally, we assign a measure of strength or frequency to each TAD based on the product of the probabilities generated by wavelet-transforms for each boundary and the loop anchor.

### WaveTAD identifies TADs independent of resolution, handles low signal-to-noise ratios, and provides robust and reproducible TAD calling

To exemplify the importance of a resolution-free TAD caller, fly (Hug et al. 2017), mouse (Lee et al. 2019) and human (Krietenstein et al. 2020) datasets were analyzed with multiple methods at different resolutions (**Supplemental Fig. S1-S3)**, These methods included TopDom, Insulation Score, IC-Finder, HiCseq, HiCExplorer, and 3DNetMod (see **Methods** for details). We first quantified the dependence on resolution by determining the number and size of called TADs (**Figure 2A** and **Supplemental Fig. S4**).

**Figure 2:**
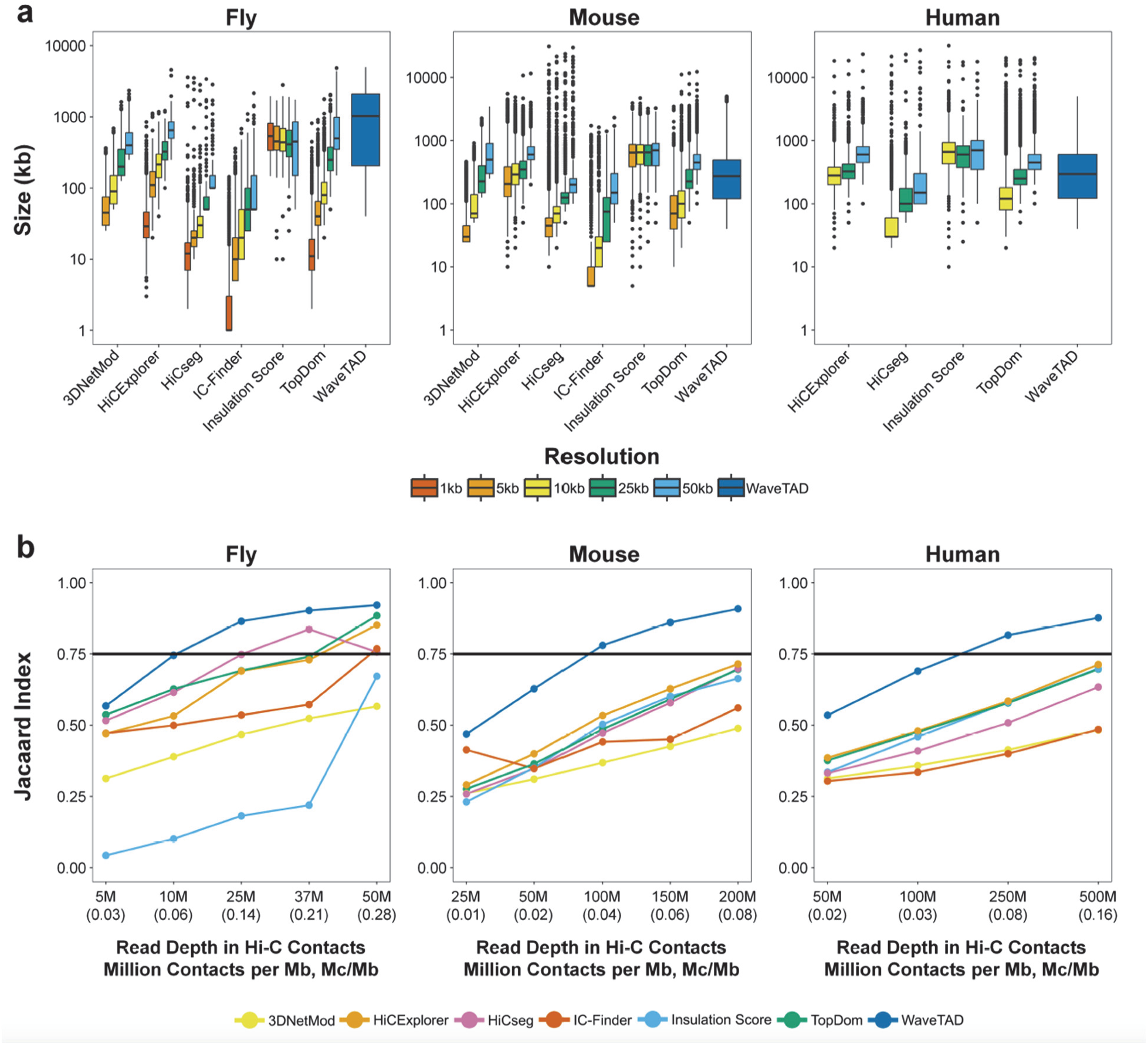
Comparison of TAD Calls between methods. **a** Box plots of TAD sizes called by each tool using different resolutions of contact matrices for each species: fly (1kb, 5kb, 10kb, 25kb, 50kb), mouse (5kb, 10kb, 25kb, 50kb), and humans (10kb, 25kb, 50kb). Note that WaveTAD is resolution-free. **b** Concordance of TAD boundaries using the Jaccard Index across read depths (number of million mapped Hi-C contacts, with million contacts per Mb in paratheses). The Jaccard Index was estimated relative to the highest contact read depth for each species: 75M (0.35 Mc/Mb) for flies, 250M (0.10 Mc/Mb) for mouse, and 1B (0.31 Mc/Mb) for human. A resolution of 10kb for fly and 25kb for mouse and human was used.

In all datasets, analyses at different resolutions drastically change the detected TADs. Finer resolutions systematically bias towards identifying more and smaller TADs, while broader resolutions reveal fewer, larger TADs. This dependency on resolution is substantial, with the number of TADs and their median size shifting more than an order of magnitude between fine and broad resolutions. On the other hand, WaveTAD calls TADs with a wide range of sizes compared to the other methods, as expected based on the identification of hierarchical 3D structures and its intrinsic resolution-free nature. Moreover, WaveTAD calls equivalent TAD sizes for the three species analyzed. Only Insulation Score calls a consistent number of TADs regardless of resolution although it calls significantly fewer TADs and with a much narrower range of sizes than WaveTAD. Because resolution in TAD studies is often different for different species and can be also influenced by the number of contact reads in a given experiment, resolution-dependent analyses of TADs could generate inappropriate information about biologically relevant properties (such as differences in average TAD size when comparing species) whereas WaveTAD allows for more direct and objective comparisons.

To explore WaveTAD’s ability to handle noisy data (low number of contact reads per Mb) better than previous methods, we subsampled the fly, mouse and human datasets. For these analyses of sensitivity and accuracy, we analyzed the boundaries called by each tool relative to those identified when using the highest read depth sample for each dataset as reference. Both the Jaccard Index (JI) (**Figure 2B)** and the overlap coefficient (**Supplemental Fig. S5)** show WaveTAD as the method least influenced by reduced read depths when compared to all other methods in the three species.

True positive rates (TPR) and false discovery rates (FDR) were also determined for each tool by comparing the subsampled reads to the highest read depth sample (**Supplemental Fig. S6**). All tools show a reduction in TPR when read depth is reduced, but WaveTAD is the only tool to maintain a high TPR while maintaining a low FDR for each read depth. Analyses of fly genomes would achieve a TPR above 0.85 with ∼25M contacts (∼0.10M contacts per Mb) with an FDR below 0.02. Analyses of mouse and human genomes would achieve TPR above 0.85 with ∼200M (∼0.08 contacts/Mb with FDR <0.05) and ∼500M (∼0.16 contacts/Mb with FDR < 0.06) Hi-C contacts, respectively. The contact matrices generated for different read depths and the corresponding TADs called by different methods were also plotted as heat maps to allow for visualization (**Supplemental Fig. S7-S9**).

To study the consistency of TAD-calling by the different tools, we analyzed multiple biological and technical replicates from fly datasets (**Supplemental Table S1**) (Hug et al. 2017) (see **Methods** for details). The concordance (JI) of boundaries called between biological replicates when using WaveTAD was 0.883 whereas it ranged between 0.105 (Insulation Score) and 0.575 (TopDom) for the other methods **(Figure 3**). Similarly, WaveTAD also had the greatest concordance among all tools to infer TADs between technical replicates (0.825), with Insulation Score and TopDom generating JI ranging between 0.313 and 0.639 (**Supplemental Fig. S10**).

**Figure 3.**
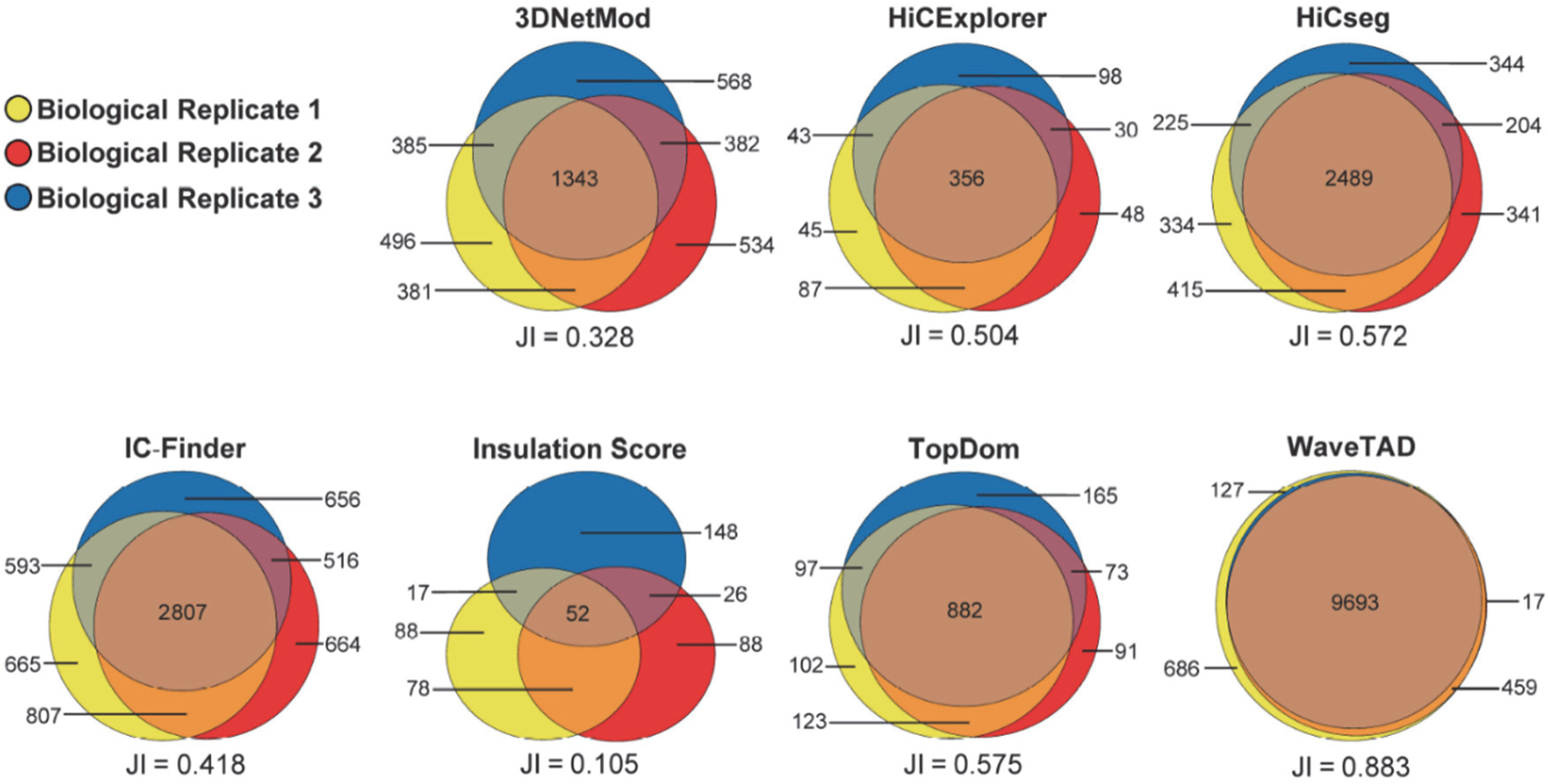
Concordance of TADs called by various tools between biological replicates in *Drosophila melanogaster*. Venn diagram depicting the number of TADs and number of overlapping TADs called between biological replicates. For each tool the Jaccard Index for all replicates is located below the Venn diagram. Data from Hug et al. staged embryos 3-4 hours post fertilization, biological replicates 1-3, (yellow, red, and blue, respectively) [30].

WaveTAD also shows a high degree of consistency when calling probabilities from biological and technical replicates. When broken up into 25 quantiles, TAD probabilities between replicates show a mean Spearman’s ρ of 0.980 and 0.972, for biological and technical replicates, respectively (see **Supplemental Fig. S11-S12** for results from 10, 25, 50 and 100 quantiles).

### Probabilities generated by WaveTAD capture TAD frequency in heterogeneous samples

To evaluate the biological relevance of the TAD probabilities produced by WaveTAD when analyzing bulk Hi-C data, we compared the frequency of a TAD being called in single-nucleus Hi-C (snHi-C) with TAD probabilities generated from bulk analysis from the same genomes. We used snHi-C from mouse embryonic stem cells (mESCs) (**Supplemental Table S1**) (Olivares-Chauvet et al. 2016; Nagano et al. 2017) and selected the top 500 single-nucleus samples in terms of read depth coverage. These snHi-C analyses allowed estimating the frequency a given TAD is called when these 500 samples are analyzed individually. We then generated an *in silico* bulk (‘merged snHi-C’) sample consisting of the first one million reads from each of the top 500 single-nucleus samples (500 million total). We also analyzed a more traditional bulk Hi-C sample from the same mESCs line (Olivares-Chauvet et al. 2016). TAD probabilities from the merged snHi-C and bulk Hi-C datasets were broken up into multiple quantiles (10, 25, 50 and 100) and compared to the frequency of the TADs called in the snHi-C dataset (**Figure 4** and **Supplemental Fig. S13**), showing high association between the frequency of TADs from the snHi-C dataset and the strength of the probability generated by WaveTAD in bulk analyses (either merged snHi-C or bulk Hi-C datasets) (e.g., Spearman’s ρ = 0.968 and 0.945, respectively, when using 25 quantiles).

**Figure 4.**
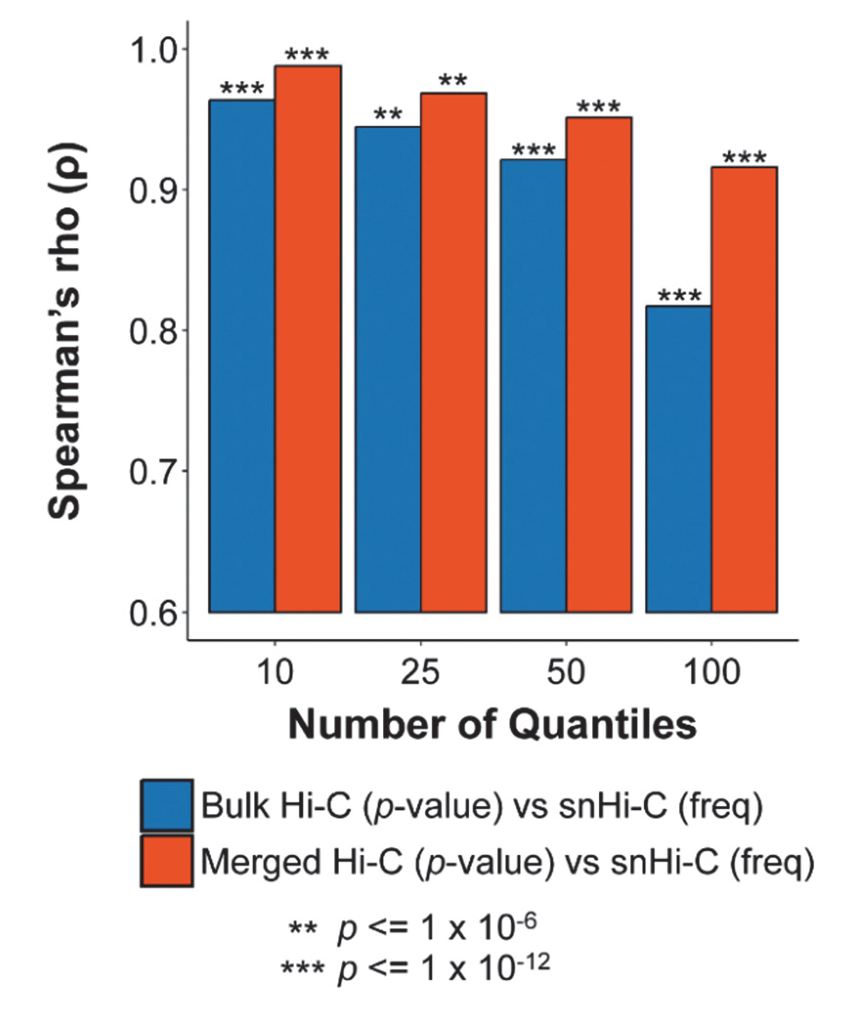
Probabilities generated by WaveTAD capture TAD frequency in heterogeneous samples. Bar plot of the correlation (Spearman’s Rank Correlation) between the frequency of a TAD being called from single-nucleus Hi-C (snHi-C) when compared to *p*-values of TADs called in merged Hi-C (blue) and bulk Hi-C (red). For each comparison, both the frequency and *p*-values of all shared TADs were placed into either corresponding 10, 25, 50, or 100 quantiles.

These results suggest that differences in probabilities associated with TADs provide information about the variable frequency of these TADs within a sample of heterogeneous genomes, thus allowing better inferences about TAD properties and dynamics with bulk Hi-C approaches.

### WaveTAD identifies TADs and boundary changes in heterogeneous samples

To examine WaveTAD’s sensitivity in calling TADs present only in a fraction of genomes, we analyzed mixed Hi-C data from two *Drosophila melanogaster* cell lines (S2 and KC167) (**Supplemental Table S1**) (Li et al. 2015; Ray et al. 2019). A sample was created to mimic a heterogenous population of cells by subsampling and mixing Hi-C contacts from the two cell lines, with either 1:0, 3:1, 1:1, 1:3, or 0:1 ratio of S2 to KC167 reads while maintaining the same total number of usable Hi-C contacts (75M). **Figures 5A-B** show the analysis of two nearby TADs called in the S2 cell line but not in the KC167 line. Both TADs are identified even when S2 Hi-C reads represent only 25% of the whole sample, with the strength of the TAD call decreasing (less statistically significant) with the fraction of genomes containing the TAD (**Figure 5B**). This property remains genome-wide when applied to all S2-specific TADs, with the median strength of these TAD calls increasing as the proportion of the TAD-containing Hi-C reads increases (**Figure 5C**).

**Figure 5.**
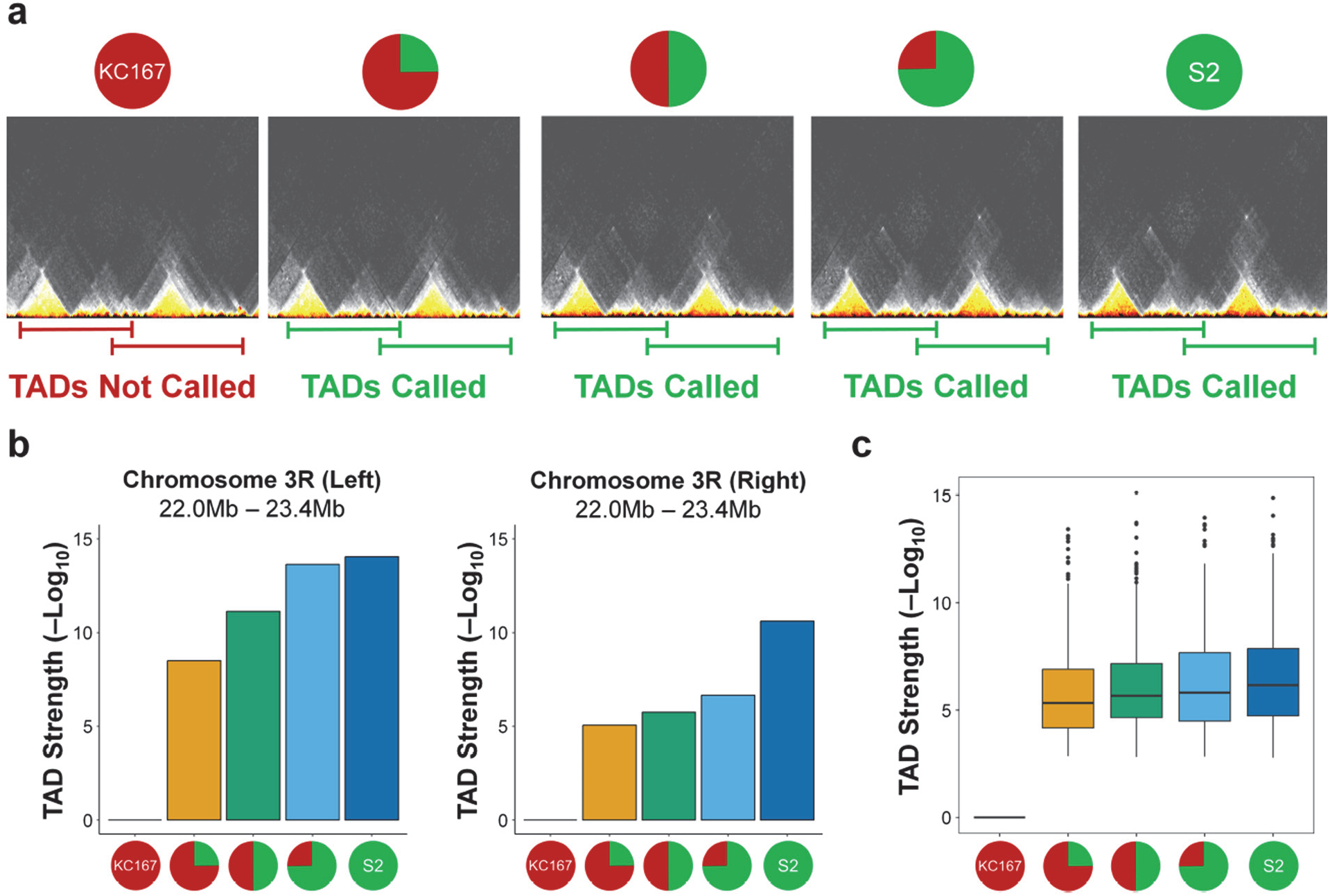
WaveTAD identifies TADs and boundary changes in heterogeneous samples. **a** Contact matrices (10kb resolution) containing Hi-C contacts mixed between two *Drosophila melanogaster* cell lines. Shown is a region (3R:22,000,000-23,400,000) with two TADs unique to the S2 cell line. The pie chart above each contact matrix shows the ratio of reads used to build the matrix (1:0, 3:1, 1:1, 1:3, or 0:1 ratio of KC167 to S2, respectively). Below, lines indicate whether WaveTAD called the pair of TADs (green) or not (red). **b** Bar plots showing the TAD strength (-Log_10_(*p*)) of each of the two TADs highlighted in **a**, across the mixed Hi-C contact samples. **c** Genome wide analysis of all TADs unique to the S2 cell line. Bar plot shows median TAD strengths (-Log_10_(*p*)) for the S2-unique TADs given the ratio of Hi-C contacts in the mixed samples (pie charts on the x-axis). For all analyses, contact matrices contained a total of 75 million Hi-C contacts.

We also assessed ability of the different methods to identify boundaries in heterozygous samples in larger genomes as a function of read depth and resolution. Specifically, we analyzed pluripotent human stem cells (hPSCs) containing a *de novo* human endogenous retrovirus subfamily H (HERV-H) insertion that creates a new TAD boundary (Zhang et al. 2019) (**Supplemental Table S1**). WaveTAD called the newly generated TAD boundaries in heterozygous for read depth ranging from 0.1M Hi-C contacts per Mb (6M contact reads) to 0.016M (1M contact reads) (**Supplemental Fig. S14**). None of the other methods was consistent in identifying the heterozygous TAD. 3DNetMod, HiCExplorer, IC-Finder and TopDom identified the boundary for some resolutions but not for others. At the lowest read depth, only 3DNetMod and TopDom correctly detected the boundary (and only at specific resolutions), while IC-Finder called a false positive. Insulation Score and HiCseg did not call the new boundary at any resolution or read depth analyzed.

### Compartment sized TADs called by WaveTAD match high-resolution imaging

To provide further support to the biological relevance of TADs called by WaveTAD and other TAD callers, we compared predicted TAD boundaries with FISH-identified long-range contacts obtained with high-resolution imaging of individual human diploid fibroblast IMR90 cells (**Supplemental Table S1**) (Wang et al. 2016). Using bulk Hi-C data from the same study, we analyzed all TADs predicted to be 500kb or longer (compartment-size structures). Three of the TAD callers identified over 80% of the FISH identified contacts, with WaveTAD (90.1%) showing the highest percentage, followed by HiCExplorer (83.5%), and Insulation Score (82.4%) (both at 50kb resolution) (**Figure 6A**). At 25kb resolution, the ability of the resolution-dependent tools to call compartments greatly deteriorated except for Insulation Score.

**Figure 6.**
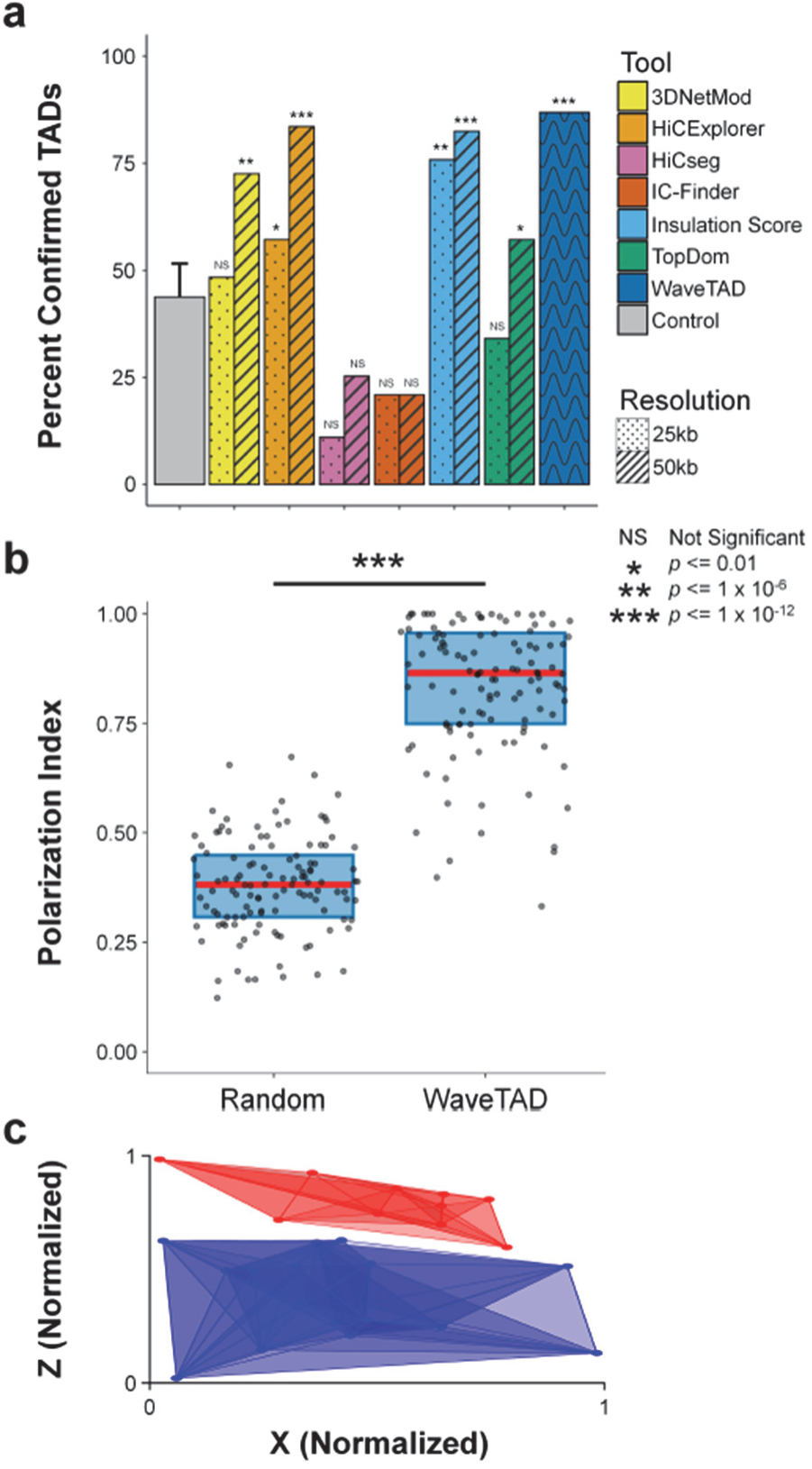
Compartment sized TADs called by WaveTAD match high-resolution imaging. **a** Percent of FISH confirmed TADS called by different tools at 25kb and 50kb resolutions (WaveTAD is independent of resolution) (N = 91, probabilities based in Permutation Test). **b** Polarization index based on the compartments confirmed by WaveTAD and a randomized control (Wilcoxon Paired Test). Analysis for chromosome 20 from FISH performed on 120 individual cells, where the red line represents the median and the blue boxes represent the 1^st^ and 3^rd^ quartiles. **c** Spatial map of the convex hulls for compartment-A (red) and compartment-B TADs (blue). Each TAD position (point) represents the centroid of the 120 individuals. The spatial map was orientated by rotating the vector connecting the centroid of the convex hulls (A and B compartments) so that it was aligned to the y-axis.

Because the genome can be divided into two compartments (A and B) that are associated with open and closed chromatin, with interactions largely constrained within the same compartment, we analyzed whether the compartment-size TADs called by WaveTAD showed separation in 3D space based on FISH data (Wang et al. 2016). We quantified the polarization index (PI) for TADs to assess overlap (or lack thereof) in 3D space, with PI expected to be 1 when compartments are completely separated in space in a polarized fashion. Large TADs called by WaveTAD are clearly divided into two different partitions, as expected if they define compartments A and B, with a median PI value of 0.865 markedly greater than the randomized control (median value of 0.381) (**Figure 6B)** that can be visually represented by building a spatial position map based on centroid positions for each of these TADs (**Figure 6C**).

### A majority of higher-order chromatin structures appear before zygotic genome activation (ZGA) in both fly and mouse

According to current views in the field, high-order chromatin structures are greatly weakened after fertilization with chromatin in a relaxed state, whereas the majority of the multi-scale 3D chromatin organization emerges alongside with zygote genome activation (ZGA) in early metazoan embryos [(Du et al. 2017; Hug et al. 2017; Ke et al. 2017); reviewed in (Jukam et al. 2017; Zheng and Xie 2019; Vallot and Tachibana 2020)]. This concept also aligns well with the key role of 3D genomic structures on gene regulation. To investigate whether TADs are indeed uncommon before ZGA or, conversely, previously missed due to weak signals that could represent transient contacts, we studied the early emergence of 3D nuclear hierarchy during fly and mouse development with published Hi-C datasets (**Supplemental Table S1**) (Du et al. 2017; Hug et al. 2017). The genome of the rapidly dividing embryos of *D. melanogaster* is initially unstructured, with a prominent surge in TAD signal occurring during zygotic genome activation (ZGA), which occurs at nuclear cycle (nc) 14 (Hug et al. 2017). Similarly, the presence of TADs becomes substantial at the 2-cell stage in mice [(Du et al. 2017; Ke et al. 2017; Zheng and Xie 2019)].

Analysis of the fly datasets using WaveTAD (**Figure 7A**) showed that the great majority of TAD boundaries present after ZGA are also identifiable before ZGA (at nc12), with a boundary concordance (JI) of 90.7%. We further compared early- and mid-stage embryo TADs to KC167 cells, a widely used and well-characterized cell line established from 6-12h embryos with a hemocyte-like mRNA expression pattern (Cherbas et al. 2011). We observed that pre-and post ZGA samples show similar boundary concordance (JI: 74.9 - 76.7%) when compared to KC167 cells. Similarly, in mice (**Figure 7B**) we observed very high concordance between boundaries called by WaveTAD at the pronuclear stage 3 (22 hours) before ZGA and inner cell masses (ICM) from blastocysts (92-94 hours) after ZGA, with a JI of 0.837. Embryonic mouse samples before and after ZGA show an equivalent high degree of TAD concordance when compared to established mouse embryonic stem cells (mESCs) (JI: 79.7 - 81.7%).

**Figure 7.**
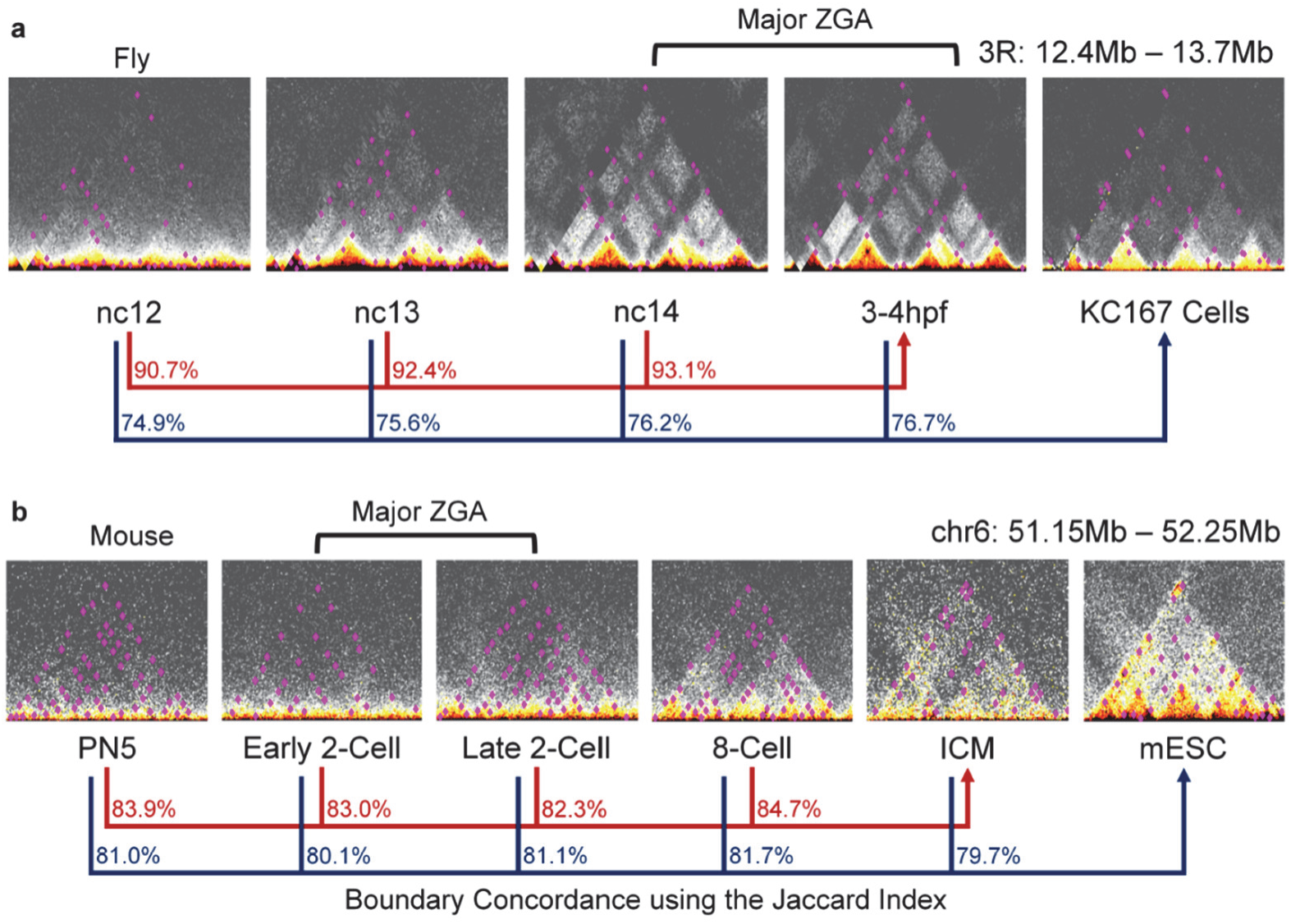
A majority of higher-order chromatin structures appear before zygotic genome activation (ZGA) in both fly and mouse. **a** Contact matrices (10kb resolution) of *Drosophila melanogaster* (chr3R:12,400,000-13,600,000) over four developmental timepoints (nc12, nc13, nc14, and 3-4hpf) and the KC167 cell line are overlayed with TADs (pink dots) called by WaveTAD. The red lines and corresponding percentages show the genome wide concordance of TAD boundaries for each timepoint (nc12, nc13 and nc14) relative to the 3-4 hpf timepoint using the Jaccard Index. The black lines and corresponding percentages show the genome wide concordance of TAD boundaries for each timepoint (nc12, nc13, nc14, and 3-4 hpf, respectively) relative to the KC167 cell line. Note that zygotic genome activation occurs at nc14. **b** Contact matrices (10kb resolution) of *Mus musculus* (chr6:51,000,000-53,000,000) over five developmental timepoints (PN5, early 2-cell, late 2-cell, 8-cell, and ICM) and the mESC cell line are overlayed with TADs (pink dots) called by WaveTAD. The red lines and corresponding percentages show the genome wide concordance of TAD boundaries for each timepoint (PN5, early 2-cell, late 2-cell, and 8-cell, respectively) relative to the ICM timepoint. The black lines and corresponding percentages show the concordance of TAD boundaries for each timepoint (PN5, early 2-cell, late 2-cell, 8-cell, and ICM, respectively) relative to the mESC cell line. Note that zygotic genome activation in mouse occurs at the 2-cell stage.

Our analyses using WaveTAD identifies pre-ZGA 3D structures that were previously overlooked likely due to widespread weaker signals, a scenario compatible with pre-ZGA contacts being highly transitory, causing a population of embryonic 3D genomes with TADs at low frequency. To seek support for this possibility and given WaveTAD’s property of generating probabilities that capture differences in frequency in heterogeneous populations, we compared the probabilities associated with TADs before and after ZGA. In both mouse and fly, probabilities associated with TADs increase in significance from before to after ZGA (Mann–Whitney U test P <2.2×10^-16^ in both fly and mouse). Thus, while WaveTAD suggests that most potential 3D genomic contacts predate ZGA and are conserved throughout development in both flies and mice, it also identifies pre-ZGA structures as less frequent—likely less stable—than those post-ZGA, supporting temporally dynamic pre-ZGA TADs. Recent super-resolution live-cell imaging inferring the existence of very transient structures in mouse embryonic stem cells would support this possibility (Gabriele et al. 2022).

### CTCF sites and TAD stability

CTCF is widely regarded as the predominant architectural protein in humans, being responsible for constraining cohesin-mediated loop extrusion via convergently oriented binding sites (Wendt et al. 2008; Phillips and Corces 2009; Alipour and Marko 2012; Ong and Corces 2014; Rao et al. 2014; de Wit et al. 2015; Nichols and Corces 2015; Sanborn et al. 2015; Dixon et al. 2016; Fudenberg et al. 2016; Dorsett 2019; Nanni et al. 2020). The 4D Nucleome (4DN) project (Dekker et al. 2017), for instance, assesses the quality of TAD calls based on the overlap with CTCF sites (Consortium 2012; Liu et al. 2015; Zhang et al. 2020) and reproducible results across different protocols and sequencing depths. The 4DN project, however, uses Insulation Score to call TADs. Given the difference in sensitivity, specificity, and reproducibility between WaveTAD and Insulation Score shown above, we re-analyzed multiple 4DN datasets of the human embryonic stem cell line H1 (H1-hESC), including standard Hi-C (Akgol Oksuz et al. 2021), 4DN Tier1 Hi-C (Krietenstein et al. 2020), and 4DN Tier1 Micro-C (Krietenstein et al. 2020), where 4DN Tier1 denotes samples with the highest sequencing depth and Micro-C identifies the highest 3D resolution protocol.

Out of the reported 69,108 CTCF sites, a much larger fraction overlap with boundaries called by WaveTAD than by Insulation Score (53.4 vs 9.7%, 71.8 vs 9.7%, and 87.3 vs 15.3% for 4DN standard Hi-C, 4DN Tier1 Hi-C, and 4DN Tier1 Micro-C, respectively). WaveTAD boundaries are also more enriched in CTCF sites than boundaries identified by Insulation score. Importantly, this is not because WaveTAD calls more TAD boundaries than Insulation Score (**Supplemental Fig. S15**). Combined, these results support the notion that WaveTAD is more sensitive and accurate than Insulation Score identifying the impact of CTCF on TADs.

Taking advantage of the quantitative information provided by WaveTAD, we then studied the impact of number and orientation of CTCF sites on TAD structures (**Figure 8**). As expected, TADs with one or more CTCF sites in at least one of the boundaries show increased statistical significance (smaller probability) than TADs without CTCF sites (*p* = 1.06 x 10^-15^), indicating improved stability and higher frequency among cells in the sample. For TAD boundaries with a single CTCF at each side, there is no difference in estimated *p*-values for convergent, divergent or same direction CTCF sites (*p* > 0.25 in all 3 pairwise comparisons). TADs with complex CTCF structures at their boundaries (multiple CTCF at each side) produce the strongest TADs of any CTCF class. This information supports the previously reported phenomena that TADs are more frequently identified when associated with multiple architectural proteins (Sexton et al. 2012; Van Bortle et al. 2012; Li et al. 2015; Cubenas-Potts et al. 2017; Rowley et al. 2017), and WaveTAD provides statistical support to the increased stability of 3D structures with increased number of CTCF sites from bulk Hi-C data.

**Figure 8.**
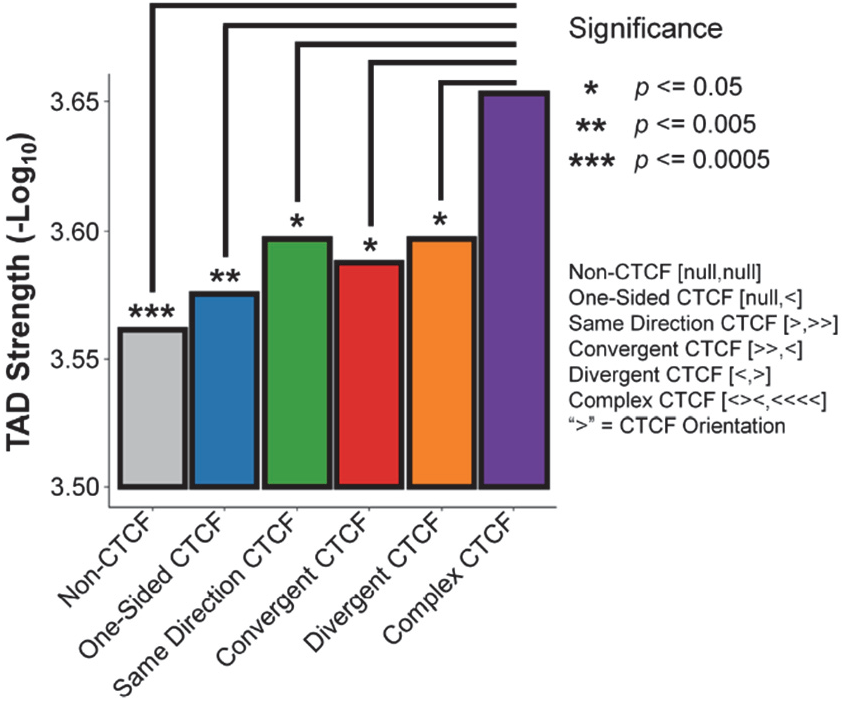
Complex CTCF sites produce more stable TADs. Bar plot showing the average TAD strength (-Log_10_(*p*)) for six classes of TAD boundaries: 1) TADs that contain no CTCF sites at either TAD boundary, 2) TADs with one boundary containing no CTCF sites and the other containing one or more CTCF sites, 3) TADs with both boundaries containing one or more CTCF sites all sharing the same orientation, 4) TADs with both boundaries containing one or more CTCF sites in a convergent orientation, 5) TADs with both boundaries containing one or more CTCF sites in a divergent orientation, and 6) TADs where each boundary contains one or more CTCF sites with at least one of the boundaries showing multiple orientations. All *p*-values were generated by bootstrapping.

### SARS-CoV-2 infection disrupts pathway-specific DNA loops

Anosmia, defined by the partial or complete loss of sense of smell, is a common symptom of SARS-CoV-2 infection and occurs in 33.9-68% of COVID-19 patients (Callender et al. 2020; Hornuss et al. 2020; Meng et al. 2020; Shamsundara and Jayalakshmi 2023). Unlike other upper respiratory infections, COVID-19 mediated olfactory dysfunction does not occur via conductive interference. Instead, a recent Hi-C and RNA-seq study of olfactory epithelium of COVID-19 patients reported downregulation of olfactory receptor (OR) and OR signaling genes together with disrupted inter-chromosomal genomic contacts, suggesting a potential disruption of the nuclear architecture in olfactory sensory neurons as the mechanism by which the virus elicits non-cell-autonomous transcriptional changes in neurons that lack entry receptors (Zazhytska et al. 2022). This study, however, only identified a significant reduction in long-range inter-chromosomal contacts (*in trans*) between clusters of olfactory receptor genes, possibly due to an elevated noise-to-signal ratio. We investigated whether WaveTAD could identify additional 3D properties using the same Hi-C and RNAseq datasets that would provide further links with pathway-specific changes in gene expression regulation (Zazhytska et al. 2022) **(Supplemental Table S1)**.

Reanalysis of RNA-seq data from olfactory epithelium of COVID-19 infected patients points out the “Olfactory Transduction” (OT) pathway as the most significantly enriched pathway, with downregulation of olfactory receptor signaling genes (**Supplemental Fig. S16**). Notably, “Apoptosis” pathway is not significantly enriched, thus ruling out cell death as a cause of differences between control and COVID-19 Hi-C samples. WaveTAD identified significant changes in intra-chromosomal structures. More specifically, this variation is driven by weakening of chromatin loops rather than boundaries, suggesting more labile promoter-enhancer interactions in COVID-19 patients (**Figure 9A-B**).

**Figure 9.**
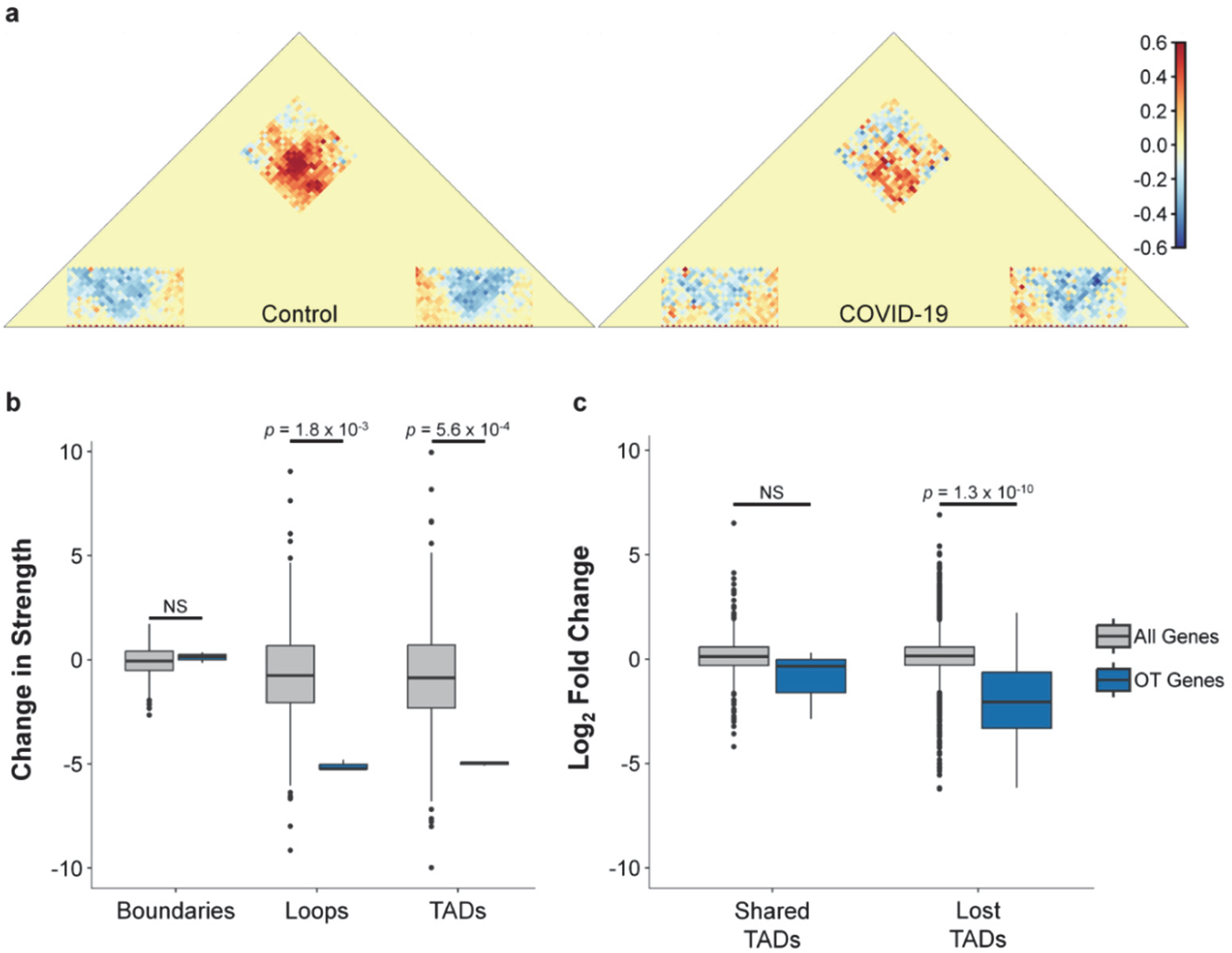
SARS-CoV-2 infection disrupts pathway-specific DNA loops. **a** Merged contact matrices of TADs containing olfactory transduction genes at one or both of their boundaries. Contact matrices were normalized to a randomized control. Areas of greater than expected contact densities are shades of red, whereas less than expected contacts densities are shades of blue. **b** Box plot of the change in strength of boundaries, loops and TADs for TADs that contain at least one olfactory transduction gene (blue). As a comparison all genome wide TADs that contain at least one gene at their boundaries are also plotted (grey). **c** Box plot of the change in gene expression levels (Log_2_-fold) of genes located at TAD boundaries for TADs that are either shared and lost between the control and COVID-19 samples. The box plot is further classified into shared and lost TADs that contain an olfactory transduction gene at one of their boundaries (blue) and TADs that contain any gene at one of their TAD boundaries (grey). All *p*-values were generated by bootstrapping.

This notion of weaker regulatory interactions is further supported by the reduction of expression of OT genes located at the TAD boundaries of TADs present in the control but disrupted in COVID-19 patients (**Figure 9C**). These results add an additional nuance to the proposed molecular mechanism of COVID-19 induced anosmia, with the preferential disruption of intrachromosomal contacts associated with the regulation of OT genes.

### Sex-specific genes and TAD stability in *Drosophila*

Sexual dimorphism can be attributed in large part to the differential expression of genes (Ellegren and Parsch 2007; Singh and Agrawal 2023). In *Drosophila* species, sex determination and sex-biased gene expression originate from genetic cascades initiated from the sex chromosomes (Yamamoto and Koganezawa 2013; Asahina 2018; Llopart et al. 2018). We took advantage of these sex-specific transcription differences to identify associated quantitative changes in 3D architecture. To this end, we used Hi-C datasets from *D. melanogaster* female and male cell lines (Li et al. 2015; Ray et al. 2019) and a list of conserved sex-specific genes across multiple *Drosophila* species (Llopart et al. 2018). Due to the small number of TADs with female-specific genes (130 out of a total of 2952), we focused on the 1313 TADs containing male-specific genes.

Overall, there is no evidence of changes in TAD presence/absence at genomic locations with male-specific genes when comparing male and female cell lines (*p* > 0.5). There are, however, quantitative differences in TAD and loop stability (**Figure 10A**). 3D structures with male-specific genes show significantly stronger signal at both TAD boundaries and loops in the male cell line relative to the female cell line, suggesting higher frequency of contacts in the male cell line relative to the female cell. Importantly, there was no statistical difference between the male and female cell lines when comparing TADs with genes known to show similar levels of gene expression in males and females (**Figure 10B**). Overall, this analysis suggests that the association between 3D nuclear structure and transcriptional programs in *D. melanogaster* is quantitative rather than qualitative, even in cases of sex-specific expression, thus explaining previous results suggesting a disconnect between TAD presence and gene expression in *Drosophila* (Ghavi-Helm et al. 2019).

**Figure 10.**
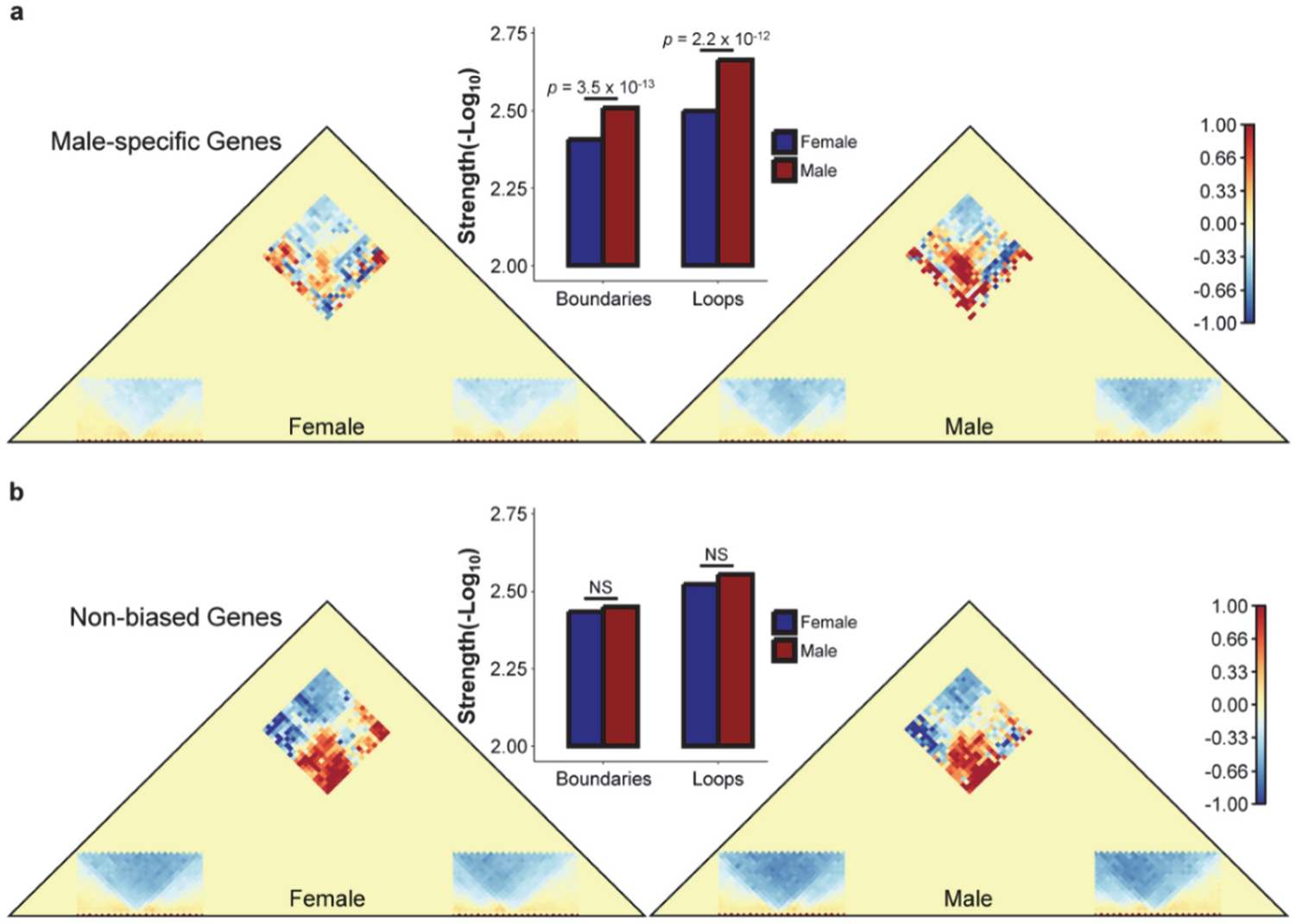
Sex-specific genes and TAD stability in *Drosophila*. **a** Merged contact matrices of TADs containing male-specific genes at one or both of their boundaries. Contact matrices were normalized to a randomized control, with greater than expected contact densities are shades of red, whereas less than expected contacts densities are shades of blue. Box plot between male and female contact matrices shows average TAD boundary and loop strengths (-Log_10_(*p*)) of TADs containing male-specific genes for the male and female cell lines (blue and red, respectively). **b** Merged contact matrices of TADs containing non-biased genes at one or both of their boundaries. Contact matrices were normalized and depicted as in **a.** Box plot between male and female contact matrices shows the average TAD boundary and loop strengths of the TADs containing non-biased genes for the male and female cell lines (blue and red, respectively).

## DISCUSSION

The accurate annotation of 3D nuclear architecture offers valuable insights. It helps us understand various cellular phenotypes related to gene regulation, development, cancer, and other diseases. Despite significant advances in experimental protocols for both bulk Hi-C and single-cell studies, current analytical frameworks are handicapped by their resolution-dependent and qualitative nature. These limitations restrict the comprehensive view of multi-layered structures, their configuration, and their dynamic nature.

Here, we described WaveTAD, a probabilistic, resolution-free, and hierarchical TAD caller based on WT. Consistent with the other WT applications, WaveTAD is better at identifying 3D structures with low signal-to-noise ratios (fewer contact reads per genomic unit) than previous methods while showing markedly higher sensitivity, specificity, and reproducibility. Whereas most of the previous methods infer a different set of loops and TADs (size, number and location) depending on the genomic unit (or resolution) used in the analyses, WaveTAD infers 3D structures without preset resolution requirements. It automatically identifies the strongest genomic signals at their optimal resolution. As such, it will allow more objective comparisons between studies of different species or when the studies differ in number of informative reads. Insulation Score, which is fairly unaffected by varying resolutions in terms of the number of TADs inferred, underperforms most other methods in terms of reproducibility, false discovery rates and the capacity to identify structures in heterogeneous samples.

The ability of WaveTAD to identify Hi-C signals in noisy datasets combined with its quantitative approach when calling structures allows detecting the presence of loops and TADs even when they are at low frequency in a sample. This property enables WaveTAD to capture differences in TAD frequency in bulk Hi-C analyses based on the probabilities associated with each structure. Until now, capturing TAD frequency has been limited to large-scale Hi-C single-cell studies but this comes at the expense of accepting limited genomic resolution. Whereas WaveTAD can be also applied to single-cell datasets (and therefore obtain frequency data directly), our studies show that it can provide both high-resolution and frequency information from Hi-C bulk heterogeneous samples.

The combination of WaveTAD new properties provides an advantage over current methods and lends a new, more quantitative approach that will be useful to study 3D dynamics and differences in probability of contact. We also demonstrated the broad applicability of this method by using different datasets and model organisms. For instance, we showed that different CTCF configurations are associated with different TAD frequencies, indicating different levels of 3D stability. We also showed that the 3D organization in early embryos is present, albeit in labile form, before ZGA in both mouse and *Drosophila* based on the study of bulk Hi-C. The comparison of TAD probabilities also allowed quantifying associations between TAD stability and differential expression in systems as disparate as COVID-19 patients or sex-specific transcription in *Drosophila*.

## METHODS

### Hi-C mapping, processing, and calculating coverage

For HiC mapping, each mate of the read pairs was mapped separately using BWA mem (Li and Durbin 2009) with parameters -E 50 -L 0 (Ramirez et al. 2018) to the reference genomes of D. emlanogaster (dm6), mouse (mm10) and human (hg38). Contact matrices were built using HiCExplorer at 1kb (flies only), 5kb, 10kb, 25kb, and 50kb resolutions (Ramirez et al. 2018). These contact matrices were then corrected using HiCExplorer with the recommended filter threshold (unique to each matrix) and normalized (Ramirez et al. 2018). If necessary for a TAD caller, Cooler dump was used to convert the contact matrices into contact tables (Abdennur and Mirny 2020).

For WaveTAD, mapped reads were filtered to only include read pairs having intrachromosomal interactions, a minimum insert length of 500bp, and a maximum insert length of 5Mb. The mates were also split based on whether it was the 5’ or 3’ mate. The coverage, in the form of a per-base report, was then calculated for the 5’ and 3’ groups using bedtools genomecov (parameters: -d) (Quinlan and Hall 2010). Regions with zero coverage depth were removed and the coverage was subsequently log transformed.

### TAD boundary calling with wavelet transforms

To generate potential TAD boundaries, wavelet-transforms were used on the 5’ and 3’ read depth coverages in a two-step approach using Maximum Overlap Discrete Wavelet-Transform (MODWT) with the coiflet filter (c6) from the R package ‘wavelets’ (Aldrich 2013). The first wavelet-transform denoises the coverage data based on the scale closest to the read pair length. The second wavelet-transform calculates the detail coefficients given by

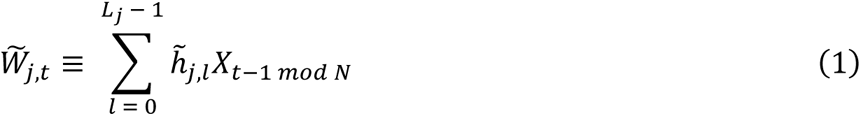

where *j* is the scale, *t* is the location, *l* is the length index, 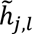 is the coiflet filter, and *X_t_*_-1_ *_mod N_* is the circulatory filter (Percival and Walden 2006). Probabilities are then derived from the detail coefficients, which are normally distributed. Assuming indepdence between adjacent data values, the variance of the detail coefficients at a given scale should be half the variance of the previous scale (Percival and Walden 2006). In genomic analyses, however, such an assumption is unlikely to be fully correct, particulalry at smaller scales. To be conservative, we chose to use the variance of the previous, smaller, *jt*ℎ scale (Equation 2).

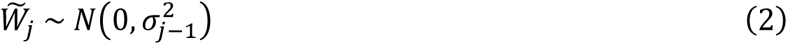

The *p*-values were then adjusted for multiple comparisons using the Holm method. Adjusted *p*-values were filtered using an α = 0.05. At locations where neighboring base pairs also passed this threshold, the mean location of the run of significant base pairs was used. Overall, the result is two lists of the potential 5’ boundaries and 3’ boundaries.

After using wavelet-transforms and its statistical framework to assign probabilities to boundaries, the potential 5’ and 3’ boundaries were paired to form potential TADs. Potential TADs smaller than the minimum TAD size (40kb in both flies and mammals (Bonev and Cavalli 2016)) and larger than a 5Mb (more lenient than the proposed 1Mb in flies (Cubenas-Potts et al. 2017) and 3Mb in mammals (Bonev and Cavalli 2016)) were removed. Potential TADs were further selected based on the presence of a loop anchor as determined by HiCExplorer’s hicDetectLoops algorithm (Wolff et al. 2020), which incorporates a donut layout proposed by HICCUPS (Rao et al. 2014; Durand et al. 2016). This algorithm analyzed the contact matrices at every possoble resolution. If a loop anchor was called at more than one resolution the smaller p-value was used downstream processes. If the coordinates of the proposed TAD failed to overlap with a loop anchor, it was removed.

In *Drosophila melanogaster*, strong TAD boundaries contain multiple architectural protein binding sites (APBS) (Cubenas-Potts et al. 2017). Therefore, the coverage across this region may contain a run of narrowly spaced spikes, which may be recognized by WaveTAD as multilpe adjacent individual boundaries. To prevent this sitation, a diamond area algorithm is applied. Some TAD callers use a variation of the algorithm first used by TopDom to call TADs (Shin et al. 2016; Ramirez et al. 2018; An et al. 2019), while others use a similar approach by comparing the interaction frequency within a proposed TAD to the surrounding frequency (Oluwadare and Cheng 2017; Roayaei Ardakany and Lonardi 2017; Chen et al. 2018; Lyu et al. 2020). We apply the TopDom algorithm at each resolution and use these boundary calls proportionately to the size of the proposed TAD being called by WaveTAD, with the remaining boundary p-value being the minimum that WaveTAD called for the individual boundaries. The result is a TAD call whose probability is the product of the loop anchor, 3’ boundary, and 5’ boundary probabilities.

### Parameters of the different TAD calling methods used for comparison

The 3D tools chosen for comparison were selected based on their diversity of approach, popularity, and third-party assessments (Dali and Blanchette 2017; Forcato et al. 2017; Zufferey et al. 2018). The algorithmic approaches included network features (3DNetMod (Norton et al. 2018)), linear score (HiCExplorer (Ramirez et al. 2018), Insulation Score (Crane et al. 2015), and TopDom (Shin et al. 2016)), statistical model (HiCseg (Levy-Leduc et al. 2014)) and clustering (IC-Finder (Haddad et al. 2017)) (Zufferey et al. 2018). Using the number of citations as the criteria for popularity, each tool is among the top for each of the algorithm classes. For all tools, the default user-defined parameters were used except where deviations were recommended by the authors or other published materials.

3DNetMod (v.1.0) (Norton et al. 2018) used parameters from the settings file when downloading the tool (settings_TEST.txt). Input matrix was a raw contact matrix. The tool did produce an unfixable error when analyzing large (high-resolution) matrices. Parameters: region_size=150, overlap=100, logged=True, badregionfilter=True, plateau=3, chaosfilter=True, chaos_pct=0.85, num_part=20, size_threshold=4, var_thresh1=0, var_thresh2=35, var_thresh3=40, var_thresh4=100, var_thresh5=0, size_s1 = 400000, size_s2 = 800000, size_s3 = 1600000, size_s4 = 3000000, size_s5 = 12000000, boundary_buffer=bin size of data.

HiCExplorer (v.2.2.1) (Ramirez et al. 2018) used all default parameters. Input was a corrected and normalized matrix.

HiCseg (v.1.1) (Levy-Leduc et al. 2014) used the same parameters from Forcato et. al. (Forcato et al. 2017) who based their parameters on suggestions from the author. Input matrix was a raw contact matrix. Parameters: size_mat=size of matrix, nb_change_max=1/3 matrix size in bins, distrib=”G”, model=”D”.

IC-Finder (v.1.0.0) (Haddad et al. 2017) used all default parameters. Input was a corrected and normalized matrix.

Insulation Score (v.1.0.0) (Crane et al. 2015) used all default parameters. Input was a corrected and normalized matrix. Online documentation noted that Insulation Score script was intended for matrices less than 10,000 by 10,000, which as noted would lead to long run-times. To avoid these run-times, the large (high-resolution) matrices were partitioned into a series of overlapping 10,000 by 10,000 matrices. This drastically brought down run-time and comparisons showed identical results (data not shown).

TopDom (v.0.8.2) (Shin et al. 2016) used the same parameters from Zefferey et. al (Zufferey et al. 2018) who based their parameters on suggestions from the author. Input was a corrected and normalized matrix. Parameters: window.size=5.

### Comparing TAD boundaries between samples

The Jaccard Index (JI) and the overlap coefficient were used as metrics to measure concordance between samples, conditions or methods. TAD boundaries were considered the same between two sets if they were within one bin of each other on either side. For WaveTAD, which calls a boundary at a single nucleotide location rather than a bin, the amount of error on each side was matched to the bin size leeway of the other tools. This meant however, that when given a large bin a single boundary could be concordant with more than one boundary. Thus, the intersection of A and B (where A is the set of TAD boundaries in one sample and B is the set of TAD boundaries in another sample) may be different from the intersection of B and A, which lead to a slight modification to the Jaccard Index and overlap coefficient equations. The Jaccard Index was defined as follows:

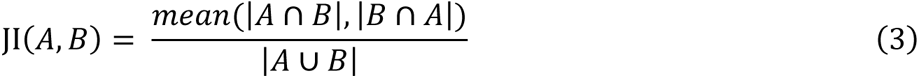

Where JI is the Jaccard Index, *A* is the set of TAD boundaries in one sample and *B* is the set of TAD boundaries in another sample. While the overlap coefficient was defined as follows:

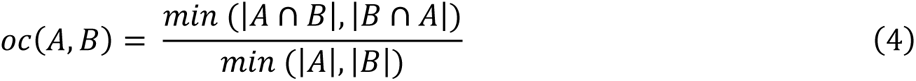

Where *oc* is the overlap coefficient, *A* is the set of TAD boundaries in one sample and *B* is the set of TAD boundaries in another sample.

### Determining true positive rate and false discovery rate

As a statistical measure of performance, true positive rates (TPR) and false discovery rates (FDR) were calculated. These metrics were used to evaluate tool performance across read depth. To do so, true positives (TP) were defined as TAD boundaries called in the highest read depth sample, false negatives (FN) were defined as TAD boundaries not called in a subsampled sample that were called in the highest read depth sample, and false positives (FP) were defined as TAD boundaries that were called in a subsampled sample but not called in the highest read depth sample. Consistent with the calculation of concordance, a cushion of one bin on each side of a called TAD boundary was implemented when determining whether calls matched, with WaveTAD getting a comparable bin based on the resolution used by the other tools. Based on these definitions, TPR and FDR were calculated using the following equations:

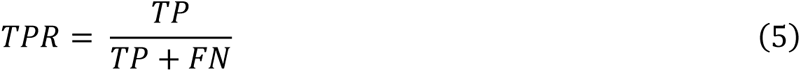

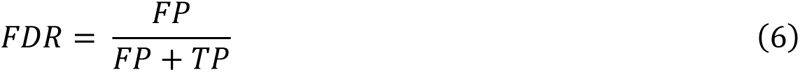

### Comparing single-nucleus Hi-C data to merged and bulk Hi-C data

The single-nucleus Hi-C dataset from mouse embryonic stem cells (see **Supplemental Table S1**) was demultiplexed following the scHiCExplorer documentation and using the scHicDemultiplex algorithm (Nagano et al. 2017; Wolff et al. 2020). The single-nucleus Hi-C dataset was then mapped, processed and analyzed by WaveTAD in the same fashion as bulk Hi-C. For the single-nucleus Hi-C dataset, only the top 500 single-nucleus samples in terms of read depth coverage were chosen and the TAD calls were combined into a single file. For the ‘merged snHi-C’ dataset, the first one million reads from each fastq file of the same 500 single-nucleus samples were appended to create the two split-read fastq files that were then treated as a bulk Hi-C sample in terms of mapping, processing and WaveTAD analysis. The ‘bulk Hi-C’ dataset represents a multi-cell Hi-C experiment using the same cell line than the single-nucleus Hi-C dataset (Olivares-Chauvet et al. 2016; Nagano et al. 2017).

After all TADs had been called for the three datasets, the TAD coordinates were converted to 50kb increments and subsequently converted to a table. Thus, each table row for the single-nucleus Hi-C data consisted of the TAD coordinates and the frequency that the TAD occurred for the 500 samples, while each table row for the merged and bulk Hi-C data consisted of the TAD coordinates and strength of the TAD (product of the loop anchor, 3’ boundary, and 5’ boundary probabilities). When comparing the samples to each other, only the intersection of the sets that had am assigned probabiity or nonzero frequency were compared. The merged and bulk TAD strengths (probabilities) were then broken down into quantiles and correlated to the inverse mean TAD frequency for each quantile from the single-nucleus Hi-C data.

### Identifying heterogeneous TADs and TAD boundaries

To mimic a heterogenous population of cells, the *Drosophila melanogaster* cell lines S2 (Ray et al. 2019) and KC167 (Li et al. 2015) were subsampled to 75 million valid Hi-C contacts and processed using the WaveTAD pipeline. TADs present in the S2 sample and absent in the KC167 sample were then identified. Mixed samples were created by mixing various ratios (3:1, 1:1 or 1:3 ratios) of KC167 and S2 contacts. TADs in the mixed samples were considered concordant to the S2 unque TADs if their respective boundaries were within 50kb.

The HERV-H insertion, and corresponding wildtype Hi-C datasets were downloaded, processed, and analyzed by the different methods to produce all TAD calls for the human chromosome 20. The precise location of the HERV-H insertion was previously documented and converted from its hg19 reference location to its hg38 counterpart. A tool’s TAD boundary call was considered a correct HERV-H insertion boundary call (true positive) if the insertion site were at most one neighboring bin away from the insertion site. A tool’s TAD boundary call was considered a false positive boundary call if they called the insertion site (under the same criteria) in the wildtype sample. This process was expanded to varying read depths (1 million, 2 million, and 3 million).

### Comparing high-resolution imaging of TADs to Hi-C derived TADs

We used high-resolution images of compartment-sized TADs from human diploid fibroblast IMR90 cells (Dixon et al. 2012; Wang et al. 2016) (**Supplemental Table S1**). These FISH images were obtained after designing probes to bind within the individual compartments (Wang et al. 2016). Subsequent analysis validated that A and B compartments are spatially organized in single chromosomes and that the spatial distance was correlated to the inverse Hi-C frequency from these same IMR90 cells. We used the IMR90 raw Hi-C dataset (Dixon et al. 2012) and mapped, processed, and analyzed it by the different TAD-calling tools. To determine whether each tool accurately calls compartment-sized TADs, all TADs under 500kb in size were first filtered out from the individual tool calls. The remaining compartment-sized TADs were then compared to the midpoint of the FISH confirmed TADs. If they overlapped with the midpoint, then the conclusion was that the called TADs overlapped with at least 50% of the image-verified TAD and we called the Hi-C inferred TAD was confirmed.

The polarization index was calculated for each FISH replicate of chromosome 21 based on the previously described compartment assignments and the following equation (Wang et al. 2016):

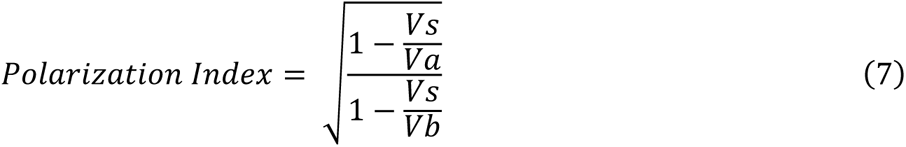

Where *Va* is the convex hull volume of compartment A TADs, *Vb* is the convex hull volume of compartment B TADs, and *Vs* is the shared volume.

To create the spatial map of the compartments, we calculated the centroid position of all the individual TADs for the 120 replicates. The A and B compartment assignments were based on the correlations between the PCA and chromatin state described in (Wang et al. 2016). We then calculated the convex hull for each compartment and their centroid position and defined the vector between each compartment’s centroid as the polarization axis. The compartments were then rotated so that the polarization axis was in the z direction.

### CTCF and TAD properties

To investigate the association between TADs and CTCF presence we took advantage of human Embryonic Stem Cell (hESC) line H1 Hi-C and Micro-C datasets (Krietenstein et al. 2020; Akgol Oksuz et al. 2021) used in the 4D Nucleome (4DN) project domain calling pipeline (Dekker et al. 2017). For CTCF sites, we used ENCODE CTCF ChIP-seq data (Consortium 2012; Liu et al. 2015), with a total of 69,108 CTCF sites reported for H1-hESC. We used previously defined CTCF orientations for these sites (Nanni et al. 2020) and calculated *p*-values using bootstrapping.

### Consistency of TAD-calling by different methods

We analyzed multiple technical and biological Hi-C replicates from Drosophila melanogaster embryos 3-4 hours post fertilization (Hug et al. 2017) (**Supplemental Table S1**) and estimated boundary concordance using the Jaccard Index (JI; see above). Note that each technical replicate contained between 29 million and 31 million valid Hi-C contacts, while the biological replicates varied between 95 million, 64 million and 68 million valid Hi-C contacts. These discrepancies in read depth likely caused some of the tools, including WaveTAD, to perform slightly better (in terms of JI) when analyzing the biological replicates rather than the technical replicates.

### Analysis of TAD strcutures in early fly and mouse embryogeneis

*Drosophila melanogaster* embryonic Hi-C datasets across four timepoints in development (nc12, nc13, nc14, 3-4hpf) were obtained from (Hug et al. 2017) (**Supplemental Table S1**). For each biological replicate, all technical replicates were combined. The result was three biological replicates for each timepoint, consisting of at least 63 million valid Hi-C contacts. The Hi-C data was then processed, and analyzed with WaveTAD. Estimates of boundary concordance using the Jaccard Index (JI; see above) for the fly dataset (**Figure 7A**) are the average JI from all possible pairwise comparisons between biogical replicates of the nc12, nc13, or nc14 timepoints with the three replicates of the 3-4hpf timepoint. All timepoint biological replicates were also compared to two biological replicates of the KC167 cell line, consisting of ∼50 million and ∼87 million valid Hi-C contacts (Li et al. 2015).

Mouse embryonic Hi-C datasets across five timepoints in development (PN5, early 2-cell, late 2-cell, 8-cell, and ICM) were obtained from (Ke et al. 2017) (**Supplemental Table S1**). All technical and biological replicates were combined to reach sufficient read depth. The result was one Hi-C sample for each timepoint consisting of at least 265 million valid Hi-C contacts.

The Hi-C data was then processed, and analyzed with WaveTAD. Estimates of boundary concordance using the Jaccard Index (JI; see above) for the mouse dataset (**Figure 7B**) are the average JI from pairwise comparisons between the PN5, early 2-cell, late 2-cell, or 8-cell timepoints with the ICM timepoint. All timepoints were also compared to the mESC cell line consisting of ∼250 million valid Hi-C contacts (Lee et al. 2019).

### Gene expression pahways in COVID-19 patients

RNA-seq (tsv files) and Hi-C datasets (pairs files) were downloaded from the 4D Nucleome Data Portal (**Supplemental Table S1**) (Dekker et al. 2017; Zazhytska et al. 2022). The RNA-seq files were directly analyzed using DESeq2 (Love et al. 2014) and subsequently used to generate a Z-score expression heatmap for olfactory transduction genes derived from a KEGG analysis. The KEGG analysis was performed using KEGGREST (Tenenbaum et al. 2019) and *p*-values obtained by a Mann-Whitney test. Hi-C pairs files were converted to sam files and analyzed by WaveTAD. Merged contact matrices were created by merging TADs (TAD size was standardized) that contained an olfactory transduction gene at one or both of its boundaries. A gene was considered overlapping if it was within 10kb of a TAD boundary. Shared TADs between control and COVID-19 samples were identified by requiring both boundaries be at most 10kb from each other. All *p*-values were generated by bootstrapping.

### Sex-specific genes and TAD stability in *Drosophila*

WaveTAD was used to identify TADs in *D. melanogaster* Hi-C datasets derived from male (S2) and female (KC167) cell lines (Li et al. 2015; Ray et al. 2019). These TADs were then filtered based on whether a gene was within 10kb of either of its boundaries. The genes were further annotated using a previously curated list of conserved sex-biased genes across multiple *Drosophila* species (Llopart et al. 2018), where each gene was categorized based on whether they were male-specific or non-biased (sex-biased and female-specific genes were removed). Merged contact matrices for male-specific genes were created by merging TADs (TAD size was standardized) that contained a male-specific genes at one or both of its boundaries in either the male or female cell line. This set of TADs was then normalized to a control set of randomized TADs. This process was repeated to create merged contact matrices for TADs containing non-biased genes. To determine whether the TAD structures composing the TADs containing male-specific genes were consistently stronger in the male cell line relative to the female cell line, a sign test was performed. As a control, this test was repeated for the non-biased genes.

## Data access

All datasets used in this study and corresponding accesion numbers are described in **Supplemental Table S1**. WaveTAD is available at https://github.com/ryanpellow84/WaveTAD

## Author contributions

J.M.C. conceived and supervised the project. R.P. and J.M.C. designed statistical and bioinformatic analyses. R.P performed statistical and bioinformatic analyses. R.P. and J.M.C. wrote the manuscript. All authors have read, revised and approved the final manuscript.

